# Distinct housing conditions reveal a major impact of adaptive immunity on the course of obesity-induced type 2 diabetes

**DOI:** 10.1101/282921

**Authors:** Julia Sbierski-Kind, Jonas Kath, Sebastian Brachs, Mathias Streitz, Matthias v. Herrath, Anja A. Kühl, Katharina Schmidt-Bleek, Knut Mai, Joachim Spranger, Hans-Dieter Volk

**Affiliations:** Department of Endocrinology & Metabolism Charité - Universitätsmedizin Berlin, corporate member of Freie Universität Berlin, Humboldt-Universität zu Berlin, Berlin, Germany; Berlin Institute of Health (BIH), Berlin, Germany; Berlin-Brandenburg Center for Regenerative Therapies (BCRT), Charité - Universitätsmedizin Berlin, corporate member of Freie Universität Berlin, Humboldt-Universität zu Berlin, Berlin, Germany; Deutsches Zentrum für Herz-Kreislauf-Forschung (DZHK), Berlin, Germany; Institute of Medical Immunology, Charité - Universitätsmedizin Berlin, corporate member of Freie Universität Berlin, Humboldt-Universität zu Berlin, Berlin, Germany; Julius Wolff Institute (JWI) and Center for Musculoskeletal Surgery, Charité - Universitätsmedizin Berlin, Berlin, Germany, corporate member of Freie Universität Berlin, Humboldt-Universität zu Berlin, Berlin, Germany; Type 1 Diabetes Center, La Jolla Institute for Allergy and Immunology, La Jolla, CA, USA; iPATH. Berlin – Core Unit Immunopathology for Experimental Models, Charité - Universitätsmedizin Berlin, Germany, corporate member of Freie Universität Berlin, Humboldt-Universität zu Berlin and Berlin Institute of Health, Berlin, Germany

**Keywords:** obesity, adaptive immunity, type 2 diabetes, non-alcoholic steatohepatitis, housing conditions, insulin resistance

## Abstract

Obesity is associated with adipose tissue inflammation, insulin resistance and the development of type 2 diabetes. However, our knowledge is mostly based on conventional murine models and promising pre-clinical studies rarely translated into successful therapies.

There is a growing awareness of the limitations of studies in laboratory mice, housed in abnormally hygienic specific pathogen-free (SPF) conditions, as relevant aspects of the human immune system remain unappreciated. Here, we assessed the impact of housing conditions on adaptive immunity and metabolic disease processes during high-fat diet. We therefore compared diet-induced obesity in SPF mice with those housed in non-SPF, so called “antigen exposed” (AE) conditions. Surprisingly, AE mice fed a high-fat diet maintained increased insulin levels to compensate for insulin resistance, which was reflected in islet hyperplasia and improved glucose tolerance compared to SPF mice. In contrast, we observed higher proportions of effector/memory T cell subsets in blood and liver of high-fat diet AE mice accompanied by the development of nonalcoholic steatohepatitis-like liver pathology. Thus, our data demonstrate the impact of housing conditions on metabolic alterations. Studies in AE mice, in which physiological microbial exposure was restored, could provide a tool for revealing therapeutic targets for immune-based interventions for type 2 diabetes patients.

## Introduction

Type 2 diabetes (T2D) is a metabolic disease that is strongly affiliated with obesity and often preceded by insulin resistance. Chronic low-grade inflammation of hypertrophic adipose tissue plays an etiologic role in the development of insulin resistance (*1*). Obesity and obesity-related complications activate both the innate and adaptive arms of the immune system and lead to the recruitment of immune cells in metabolic organ systems (*2-12*).

Currently, our understanding of the underlying mechanisms in these conditions is mainly based on experiments carried out with laboratory mice housed under standard specific pathogen-free (SPF) conditions. Those barrier facilities are unnaturally hygienic and do not reflect the microbial diversity humans encounter in their environment. Despite success in efficient translation of animal studies to the clinic, many murine models failed to translate promising treatments of disease models to clinical studies, as relevant aspects of the human immune system are unappreciated by SPF mice (*13, 14*). Recent studies claimed that murine models housed under standard SPF conditions lack effector/memory T cells in contrast to mice exposed to antigens, such as pet shop and free-living mice (*15-18*).

Thus, chronic exposure to antigens from environmental germs under “natural conditions” results in reversed proportions of naïve and effector/memory T cells and a more restricted T cell receptor repertoire, associated with changes in the B cell and myeloid compartment. This process is called “aging” of the adaptive immune system, indicated by increased frequencies of effector/memory T cells in blood and immune organs. In contrast to naïve and early memory T cells, effector/memory T cells express an altered homing and stimulation pattern enabling them to infiltrate inflamed tissue sites and contributing to intratissue inflammatory state with major impact on tissue homeostasis. Recently, we could establish a model of antigen exposure under standardized non-SPF housing conditions (*15*). Applying this model, we aimed to assess the impact of adaptive immunity over the course of obesity-related T2D. For this purpose, we analyzed the immune cell composition in blood, adipose tissue and liver as well as markers of glucose intolerance and insulin secretion in SPF and non-SPF mice [called “antigen-exposed” (AE), here], receiving normal diet (ND) or high-fat diet (HFD).

## Materials and Methods

### Animals and Diet

This study was carried out in accordance with the Guide for the Care and Use of Laboratory Animals of the National Institutes of Health and the Animal Welfare Act under the supervision of our institutional Animal Care and Use Committee. Animal protocols were conducted according to institutional ethical guidelines of the Charité Berlin, Germany, and were approved by the Landesamt für Gesundheit und Soziales (approval number G 0138/14, LAGeSo Berlin, Germany) and comply with the ARRIVE guidelines.

Mice were housed in SPF conditions or transferred to the AE animal facility at the age of 4 weeks. Supplementary Table 1 provides an overview of the specific SPF and AE housing conditions. AE housing was complemented with daily non-specific microbial exposure to antigens from bedding of mammalian laboratory animals (sheep, pigs) housed in rooms next to the laboratory mice. The exposure was guaranteed by daily handling of the laboratory animals and antigens were distributed by air passage, as the laboratory staff switched rooms on a regular basis. A high-fat diet (HFD) (60kJ% from fat, 19kJ% from proteins and 21kJ% from carbohydrates obtained from SSNIFF (E15741-34, Soest, DE)) for mice was started at 5 weeks of age and continued for either 7 or 15 weeks. Age-matched animals were maintained on ND (SSNIFF, V1534-300, Soest, DE) which provides 9kJ% from fat, 33%kJ from proteins and 58%kJ from carbohydrates. Body weight was followed weekly throughout the course of the experiment.

### Analysis of metabolic parameters

Intraperitoneal glucose tolerance test (IPGTT) was performed in overnight fasted mice by injecting glucose i.p. (1g/kg body weight; B. Braun Melsungen, Melsungen, Germany) and blood glucose levels were monitored at 0,15,30,60,120 min postglucose administration and plotted against time curves to determine the glucose tolerance. Serum insulin was measured by ELISA kit using a rat insulin standard (Crystal Chem, Downers Grove, IL). Body composition was determined by nuclear magnetic resonance using a Bruker Minispec instrument (Bruker, Woodlands, TX). Real-time metabolic analyses were conducted using a combined indirect calorimetry system (TSE Systems, GmbH, Bad Homburg, Germany) as previously described (*19*).

### Histological Analysis

Following perfusion with saline to minimize contamination with cells from the vasculature, liver and pancreatic tissue were dissected from the surrounding tissues, fixed in formalin and embedded in paraffin. The sections (4µm) were deparaffinized with xylene and rehydrated through an ethanol gradient.

For IHC staining in pancreatic sections antigen was retrieved by citric acid buffer (pH6.0) and slides were blocked with normal goat serum and incubated with anti-insulin antibody (1:200) overnight at 4°C. Slides were then washed and incubated with an HRP-conjugated secondary antibody for 1hr at room temperature. After washing in PBS HRP was visualized by NovaRED (VECTOR NovaRED Peroxidase Substrate Kit) and nuclei were counterstained with haematoxylin. Insulin-positive β cells mark islets. The average islet area and number of islets were calculated per total pancreatic area in a minimum of five sections per sample, 200µm apart. Liver was processed for routine haematoxylin/eosin and histochemical staining. Tissue collagen desposition was detected by applying the following histochemical staining protocol: The slides were incubated with a 0.1% Sirius Red solution dissolved in saturated picric acid for 1 hour washed in acidified water (0.5% hydrogen chloride), dehydrated and mounted with DPX mounting. Collagen and non-collagen components were red- and orange-stained. For IHC staining antigens were unmasked by boiling in citrate buffer for 30 minutes. Slides were blocked using 0.3% H2O2/H2O for 10 min and incubated overnight with purified monoclonal rat IgG2b kappa anti-mouse I-A/I-E 1:50. Biotinpolyclonal goat anti-rat 1:50 was used as second antibody. Slides were then incubated with Avidin-HRP (1:2000-dilution) for 30 minutes. HRP was detected with NovaRed (SK-4800) and slides were counterstained with haematoxylin/eosin.

The slides were analyzed with a Zeiss Axioplan light microscope and the images were acquired using AxioVert.

### Tissue cell preparation

We sacrificed the mice after general anesthesia. For isolation from stromal vascular cells, we removed the epididymal adipose tissues and minced them into small pieces.

We then incubated the adipose tissue in a rotational shaker (200rpm) at 37°C for 20min in collagenase solution (1mg/ml of collagenase type1 (Worthington) in PBS supplemented with 0.5%BSA and 10 mM CaCl_2_). We passed the digested tissue through a 100µm mesh and centrifuged the cell suspension at 500g for 10min. We resuspended the resultant pellet containing the stromal vascular fraction into erythrocyte lysis buffer and centrifuged again at 500g for 5min. We washed the cells again in PBS supplemented with 2% FCS and finally resuspended them in 250µl PBS supplemented with 2% FBS. We then counted the cells based on trypan blue exclusion.

### Flow cytometry

We incubated these isolated cells with the below mentioned antibodies for 20min. after applying an Fc block with CD16/32 antibody (Biolegend, 101301) for 10min. Cells were then washed twice and filtered through a 35µm mesh. The antibodies for surface staining can be found in Supplementary Table 3. The FACS analysis was performed on BD-LSR Fortessa. FACS data were analyzed by post collection compensation using FlowJo 10.0.8 (Tree Star, Ashland, OR) software.

### Participants

Blood samples were collected from a total of 21 obese women (mean ±SD, body mass index [BMI] 34.61±3.76kg/m^2^) included in a study performed as part of a larger study focusing on muscle mass regulation “Effects of negative energy balance on muscle mass regulation” (registered at https://clinicaltrials.gov, NCT01105143) at the Department of Endocrinology of the Charité University Medicine Berlin, Germany.

This study was carried out in accordance with the recommendations of the International Conference on Harmonization Guidelines for Good Clinical Practice and the Declaration of Helsinki. The protocol of the study was approved by the local Ethics Committee of the Charité - Universitätsmedizin Berlin (EA2/050/10). All subjects gave written informed consent in accordance with the Declaration of Helsinki before participating in this study.

### Human Antibody Panel

Fluorochrome-conjugated anti-human monoclonal antibodies were obtained from Beckman Coulter (Marseille, France), except anti-BDCA-2 and anti-BDCA-3, which were obtained from Miltenyi Biotec (Bergisch Gladbach, Germany). Six panel matrices for 7- to 9-fluorochrome channels defined and validated by the ONE Study consortium were used based on published results (*20*).

### Human Leukocyte Staining

For staining 100 μL of anticoagulated peripheral blood cells were stained with surface antibodies as previously described (*17*). All samples were measured on 10 color, 3 laser Navios flow cytometers (Beckman Coulter) and acquired data files were analyzed using the Kaluza software, version 1.2 (Beckman Coulter).

### Statistical analysis

The results are shown as the mean ±SEM. All analyses were performed using GraphPad Prism version 6 (GraphPad Software, San Diego, CA). A P value of <0.05 was considered significant. Depending on the distribution of data, Pearson simple coefficient or Spearman rank correlation coefficient with two-tailed significance test were used for correlation analysis, and Mann-Whitney *U* test, Student *t* test or one way ANOVA followed by Tukey’s comparison test were applied to estimate differences between groups as appropriate. The body weight, glucose and insulin time course data were analyzed by repeated measurements two way ANOVA followed by post-hoc (Bonferroni) test.

## Results

### SPF laboratory mice lack effector memory T cell subsets

To visualize differences in immune responses, we compared T cell distribution between mice and humans. Relative to adult humans (Fig. 1A), SPF mice showed lower frequencies of effector memory T cells (< 30%), near absence of effector T cells (< 5-10%), and reciprocally, higher frequencies of naïve T cells (> 60%) in blood, particularly in the CD8^+^ T cell compartment (Fig. 1B). In contrast, decreased proportions of naïve T cells (< 40%) and enhanced levels of effector memory T cells (> 25%) were observed in AE mice, which appeared much closer to the human phenotype (Fig. 1B).

**Figure 1.**
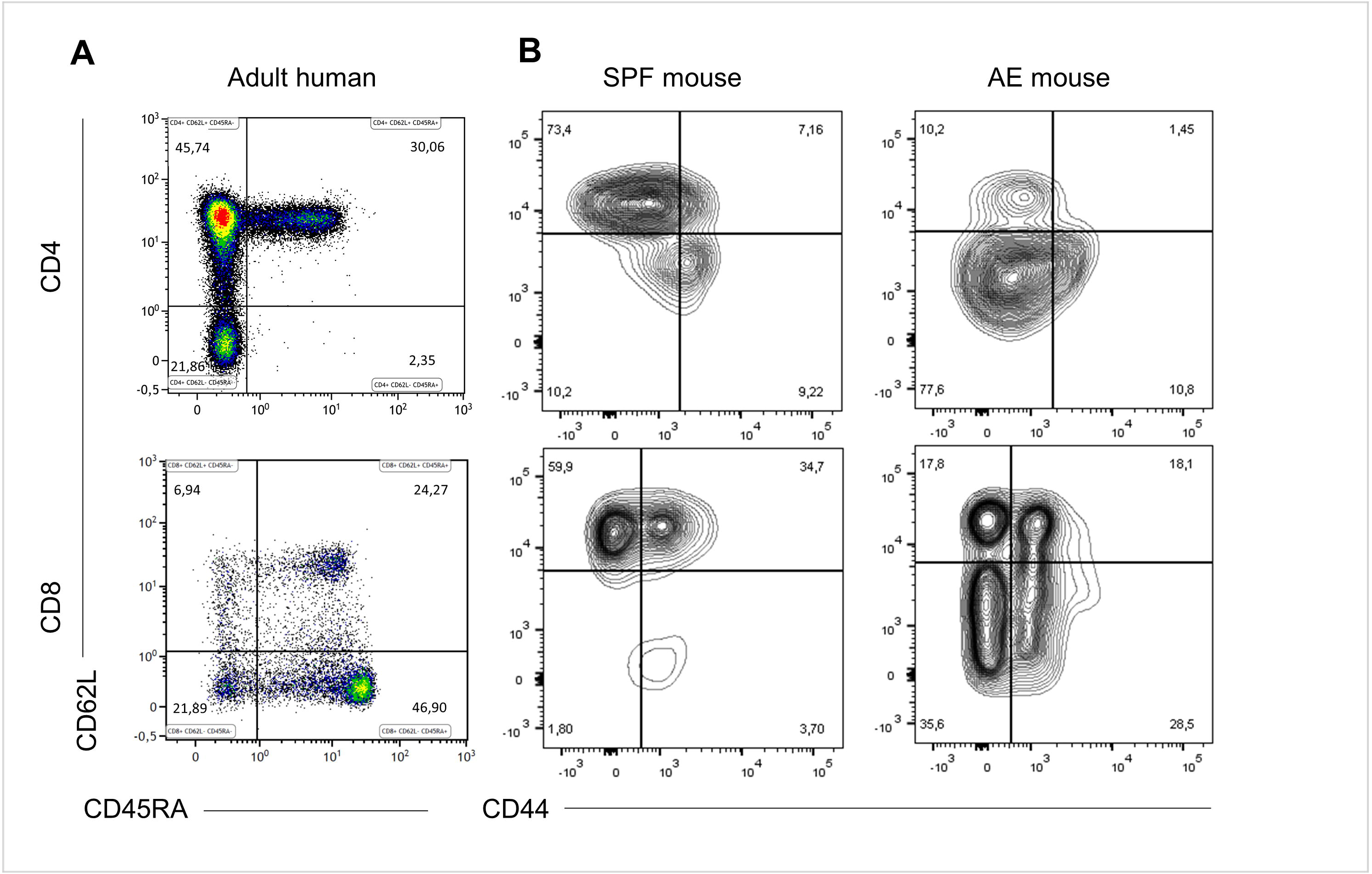
Specific pathogen-free (SPF) mice lack effector memory T cell subsets. CD4^+^ and CD8^+^ T cell phenotypes were compared between 20 weeks old SPF mouse blood (n=10), 20 weeks old antigen exposed (AE) mouse blood (n=10) and adult human blood (n=21) by fluorescence flow cytometry. Top panels are gated on CD3^+^CD4^+^ cells and bottom panels are gated on CD3^+^CD8^+^ cells and show naïve (CD44^-^CD62L^+^/CD45RA^+^CD62L^+^), central memory (CD44^+^CD62L^+^/CD45RA^-^ CD62L^+^), effector memory (CD44^+^CD62L^-^/CD45RA^-^CD62L^-)^ and effector (CD44^-^CD62L^-^/CD45RA^+^CD62L^-^) T cells. **(A)** Representative dot plots of blood T cell subset distribution in SPF mouse and adult human. **(B)** Representative dot plots of blood T cell subset distribution in AE mouse. Numbers in panels indicate percentages of T cell subsets.

### Altered immune cell composition in AE mice kept on high-fat diet

To assess the combined impact of housing conditions and diet on the immune composition as well as on the metabolic status, we compared AE and SPF mice kept on HFD or ND. The experimental design is shown in Fig. 2. All groups were metabolically characterized and blood was analyzed either at week 7 or at week 15. Similar to previous observations blood from AE mice was enriched in effector memory T cells and, correspondingly, showed lower percentages of naïve T cells (both CD4^+^ and CD8^+^) compared to SPF mice (Fig. 3A-C). Additionally, a subset of CD44^-^CD62L^-^ T cells strongly increased under AE conditions and is believed to be an equivalent to effector T cells (Temra) in humans. Remarkably, the frequency of effector memory T cells increased further under HFD in AE mice, but not in SPF mice (Fig. 3 B-C). The significantly enhanced expression of PD-1 (CD 279), associated with chronic T cell activation, in AE mice matched these data (Fig. 3D). Furthermore, the total amount of leucocytes and T cells was higher in AE mice compared to SPF mice (data not shown). Taken together, housing conditions and diet have a major impact on T cell subset distributions.

**Figure 2.**
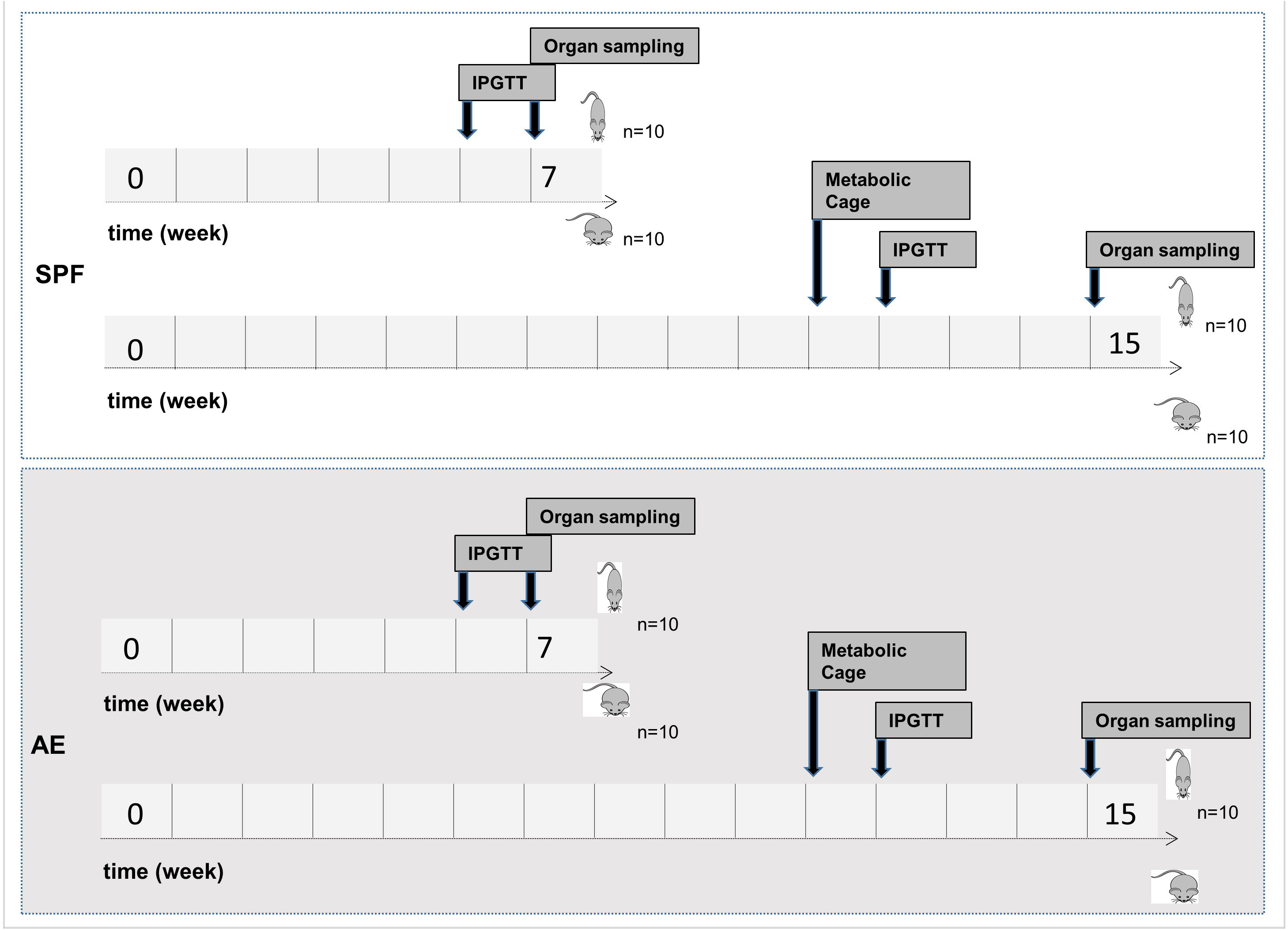
Study flowchart. Mice were kept in SPF conditions or transferred to the non-SPF animal facility at the age of 4 weeks. At the age of 6 weeks, male C57Bl/6J mice were fed either a ND or a HFD for either 7 or 15 weeks with 10 mice per group. IPGTT was performed after 6 or 12 weeks of ND or HFD feeding. Metabolic phenotyping was performed after 11 weeks of ND or HFD feeding. Animals were sacrificed at the age of 12 or 20 weeks for sample collection.

**Figure 3.**
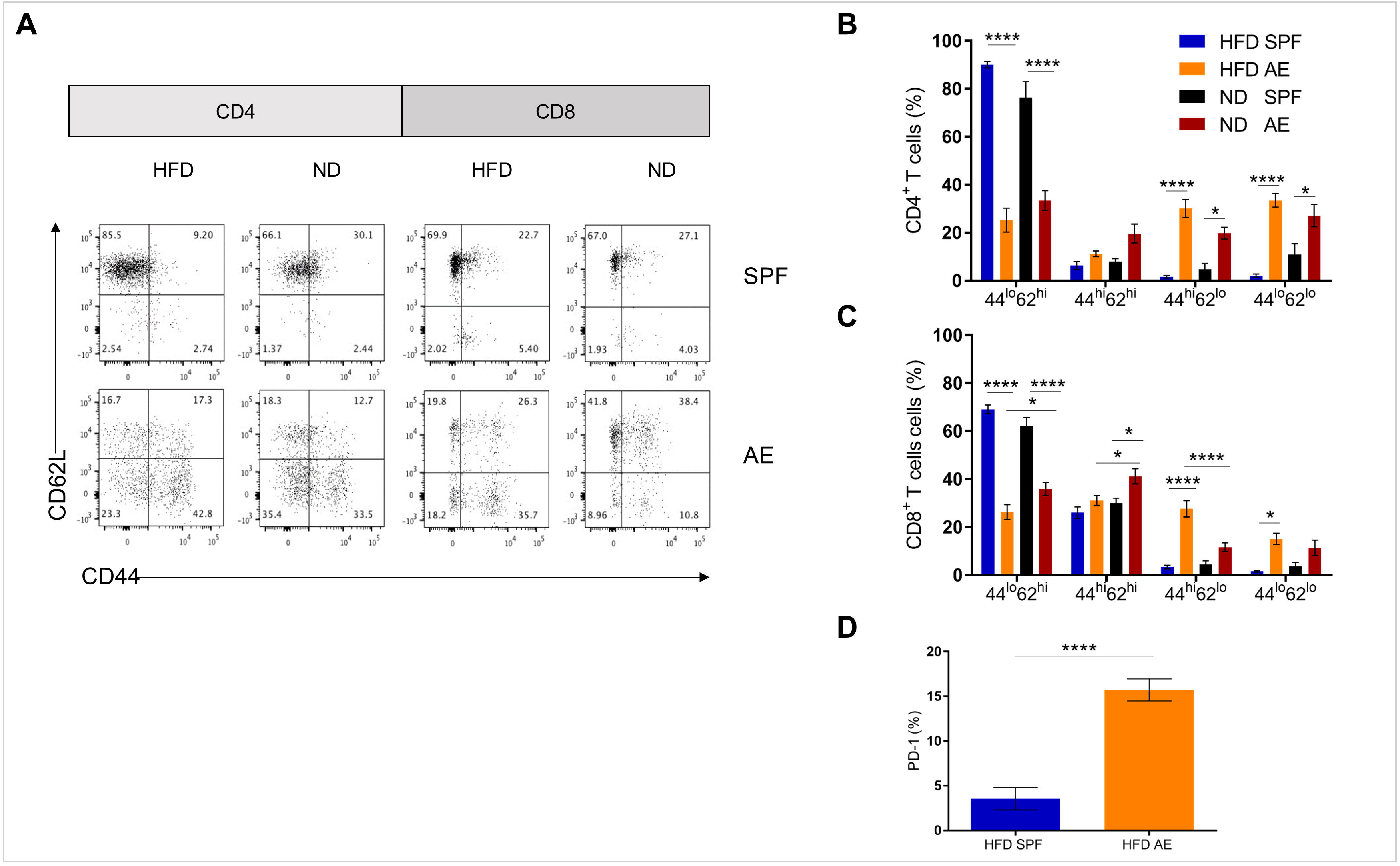
Altered immune cell composition in AE mice kept on high-fat diet**. (A)** Representative dot plots for CD4^+^and CD8^+^ T cell subsets in blood from SPF and AE mice fed a HFD or ND for 7 weeks. **(B), (C)** Percentages of CD44^lo^CD62^hi^ (naïve), CD44^hi^CD62L^hi^ (central memory), CD44^hi^CD62L^lo^ (effector memory) and CD44^lo^CD62L^lo^ (probably effector) CD4^+^ and CD8^+^ T cells. Data are represented as means ± SE. Significance was determined using 2-way ANOVA multiple measurement. **(D)** PD-1 expression of CD44^hi^CD62L^lo^T cells in HFD SPF compared to AE mice. Data are represented as means +SE. Significance was determined using unpaired 2-sided Mann-Whitney U-Test. n=10 mice per group. *P<0.05, *** P<0.001,****P<0.0001.

### Preserved pancreatic ß-cell responsiveness in HFD AE mice

To assess the impact of “aged” adaptive immunity following antigen exposure on the development of insulin resistance and glucose intolerance, we examined the metabolic effects of HFD long-term feeding in SPF and AE mice. HFD feeding resulted in the development of metabolic alterations already after a period of 7 weeks in SPF mice, as shown in Supplementary Fig. 1 A-G. Within both cohorts (SPF and AE) a significant difference in body weight between HFD and ND mice was first evident at week 4 (*P* < 0.001) and remained significant throughout the following experimental weeks (Fig. 4A). HFD fed AE mice tended to gain more weight after 7 weeks and were even significantly heavier than their SPF counterparts after 15 weeks of HFD feeding. In contrast, epididymal fat pads were enlarged in HFD mice but remained unaffected by housing conditions (Fig. 4B). Whereas glucose tolerance was impaired in HFD mice maintained under SPF conditions, surprisingly, the glucose levels in AE mice did not differ significantly from those of the ND mice after 7 weeks of HFD (Fig. 4C). This striking difference was not observed at a later time point (15 weeks) in HFD fed mice although fasting and peak glucose levels remained slightly lower in AE mice compared to SPF mice (Fig. 4C). Insulin levels, measured throughout the IPGTT, were significantly higher in HFD AE mice (Fig. 4D). The insulinogenic index, the ratio of the increment in insulin concentration to the increment in glucose concentration (ΔI/ΔG) has been proposed as a measure for ß-cell function and is highest in HFD fed AE mice at all time points (Fig. 4E). In summary, HFD AE mice develop obesity and insulin resistance but show adequate ß-cell compensation and do not develop diabetes over a period of at least 7 weeks in contrast do HFD SPF mice.

**Figure 4.**
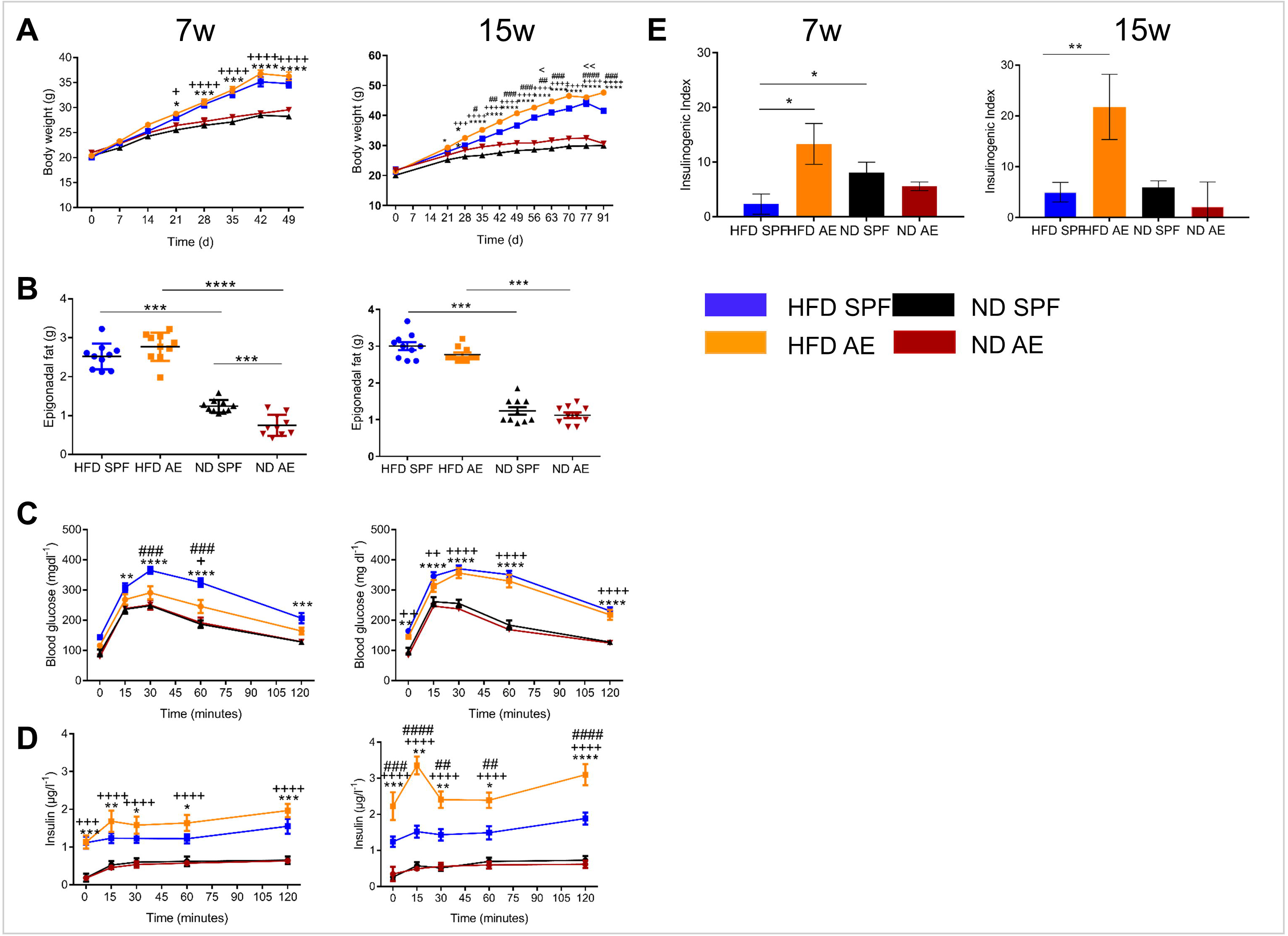
Preserved pancreatic ß-cell responsiveness in AE HFD mice**. (A)** Weight development in 7week (left) and 15week (right) ND and HFD SPF vs. AE mice. **(B)** Weight of epididymal fat pads. **(C)** Intraperitoneal glucose tolerance tests (IPGTT) were performed in ND and HFD SPF vs. AE mice after 7 weeks (left) and 15 weeks (right) feeding. **(D)** Circulating insulin levels were assessed before and after intraperitoneal glucose injection in ND and HFD SPF vs. AE mice after 7 weeks (left) and 15 weeks (right) feeding. **(E)** Calculated insulinogenic index, n=10 mice per group. Significance was determined using 2-way ANOVA multiple measurement **(A**,**C**,**D)** or using unpaired 2-sided Mann-Whitney U-Test**. (B, E)** * P<0.05, ** P<0.01, *** P<0.001, **** P<0.0001. *(HFD SPF vs. ND SPF), ++ (HFD AE vs. ND AE), # (HFD SPF vs. HFD AE).

### Diet-induced obesity rather than housing conditions change the immune cell composition in visceral adipose tissue

To further establish to what extent housing conditions and diet influence the adipose intratissue immnune status, we analyzed visceral adipose tissue by flow cytometry. Fig. 5A summarizes the relative distribution of various immune cell subsets in HFD AE and SPF mice at week 7 and 15 as a heatmap demonstrating a clear clustering of the mice.

**Figure 5.**
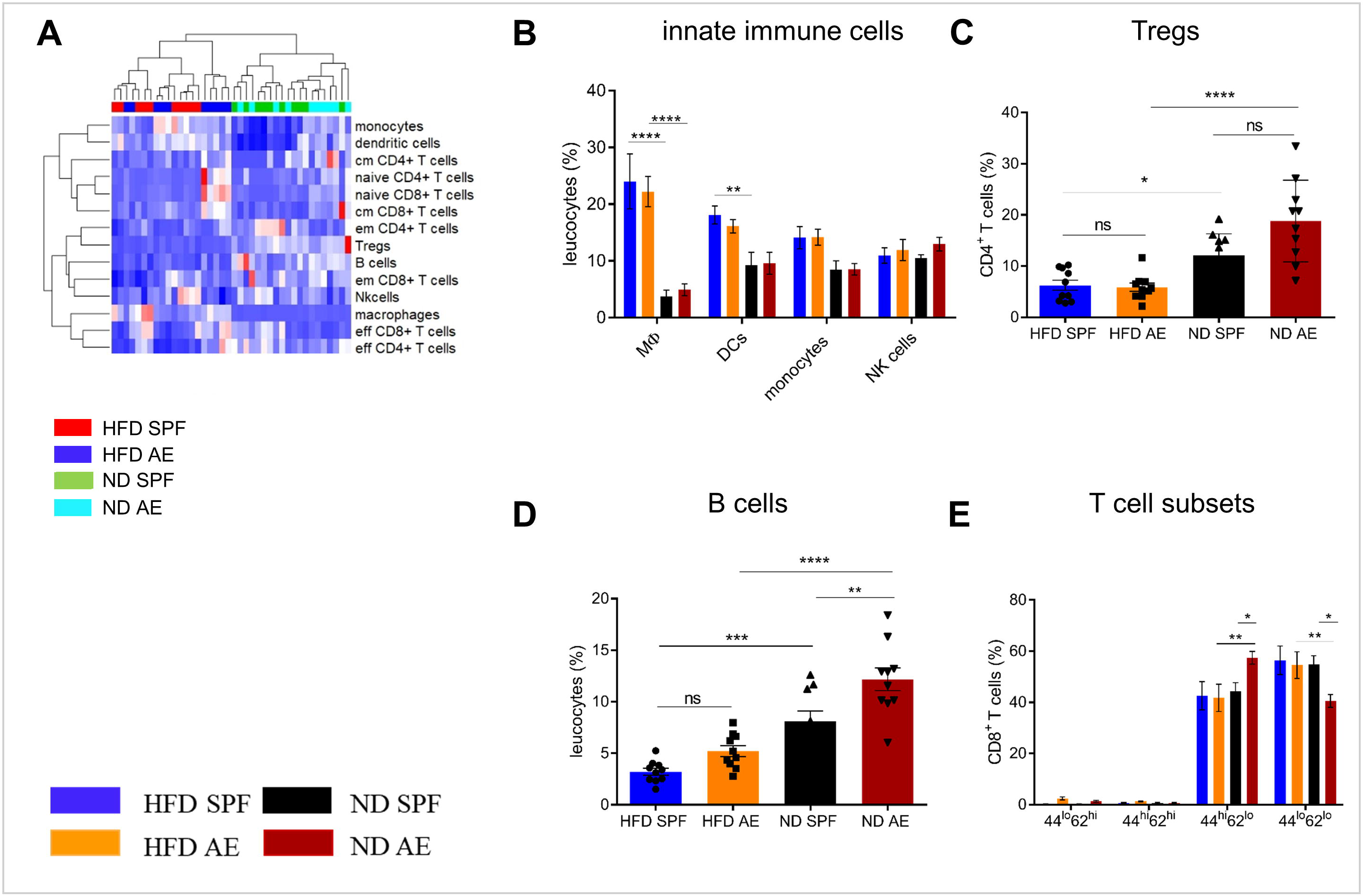
Diet-induced obesity rather than housing conditions change the immune cell composition in visceral adipose tissue. Immune cell composition was determined in visceral adipose tissue via flow cytometry in 15 weeks fed mice. **(A)** Heatmap showing relative distribution of T cell subsets and cells of the innate immune system as percentages of leucocytes in 15 week groups. Red indicates higher proportion, blue indicates lower proportion. **(B)** Distribution of innate immune cells as percentage of leucocytes in HFD and ND mice maintained under SPF or AE conditions. **(C), (D)** Proportion of regulatory T and B cells in visceral adipose tissue of SPF or AE mice. **(E)** Percentages of CD44^lo^CD62L^hi^ (naïve), CD44^hi^CD62L^hi^ (central memory), CD44^hi^CD62L^hi^ (effector memory) and CD44^lo^CD62L^lo^ (probably effector) T cells. Significance was determined using 2-way ANOVA multiple measurement **(B, E)** or ANOVA followed by Tukey’s multiple comparisons test **(C, D)** n=10 mice per group. *P<0.05, ** P<0.01, *** P<0.001, **** P<0.0001.

The percentages of macrophages (F4/80^+^CD11b^+^) and dendritic cells (CD11b^+^ CD11c^+^) were significantly higher in HFD compared to ND mice, which is in line with previous results (*21*). However, innate immune cells were equivalent between both AE and SPF cohorts of mice (Fig. 5B). Only some temporary differences in dendritic and NK cells were observed between AE and SPF mice (Supplementary Fig. 2B). Similarly, both HFD groups showed comparably decreased proportions of regulatory CD4^+^ T cells (Treg) and B cells in relation to ND mice (Fig. 5C and D, Supplementary Fig. 2C-D). As expected, in all four groups (AE vs. SPF; HFD vs. ND) mainly effector/memory T cells accumulated in the adipose tissue (Fig. 5E). Interestingly, at week 7, we could also observe naïve-like and central memory T cells in the adipose tissue from AE but not SPF mice (Supplementary Fig. 2E).

In summary, HFD fed mice housed in SPF or AE conditions displayed higher levels of distinct innate immune cells, whereas only temporary and minor differences were observed between SPF and AE cohorts.

### Antigen exposure protects ß-cell morphology and function allowing lasting compensation of insulin resistance

Next, we examined the histology of the pancreas via haematoxylin/eosin (HE) and immunohistochemical (IHC) insulin and T cell staining to understand the metabolic differences between the groups as described above. In comparison to ND mice, HFD mice developed a small degree of islet hyperplasia as described before (*22*). Furthermore, HFD AE mice showed a higher mean cross-sectional area of the pancreatic islets at week 15 compared to HFD SPF mice (Fig. 6A). Mean islet area levels were comparable in mice under SPF conditions, whereas HFD AE mice displayed higher mean islet area levels compared to ND AE mice (Fig. 6B). Higher plasma insulin levels significantly correlated with the mean islet area (Fig. 6C) in 7 and 15 week HFD mice in both the SPF and AE cohort. In the HFD SPF group grossly diminished numbers of insulin staining ß-cells were evident, while HFD AE mice showed preserved insulin staining (Fig. 6D). Notably, periductal pancreatic inflammation was neither observed in SPF nor in AE mice. Moreover, immunostaining revealed that almost all CD3^+^ cells remain in the vessels without any alterations between different groups (data not shown).

**Figure 6.**
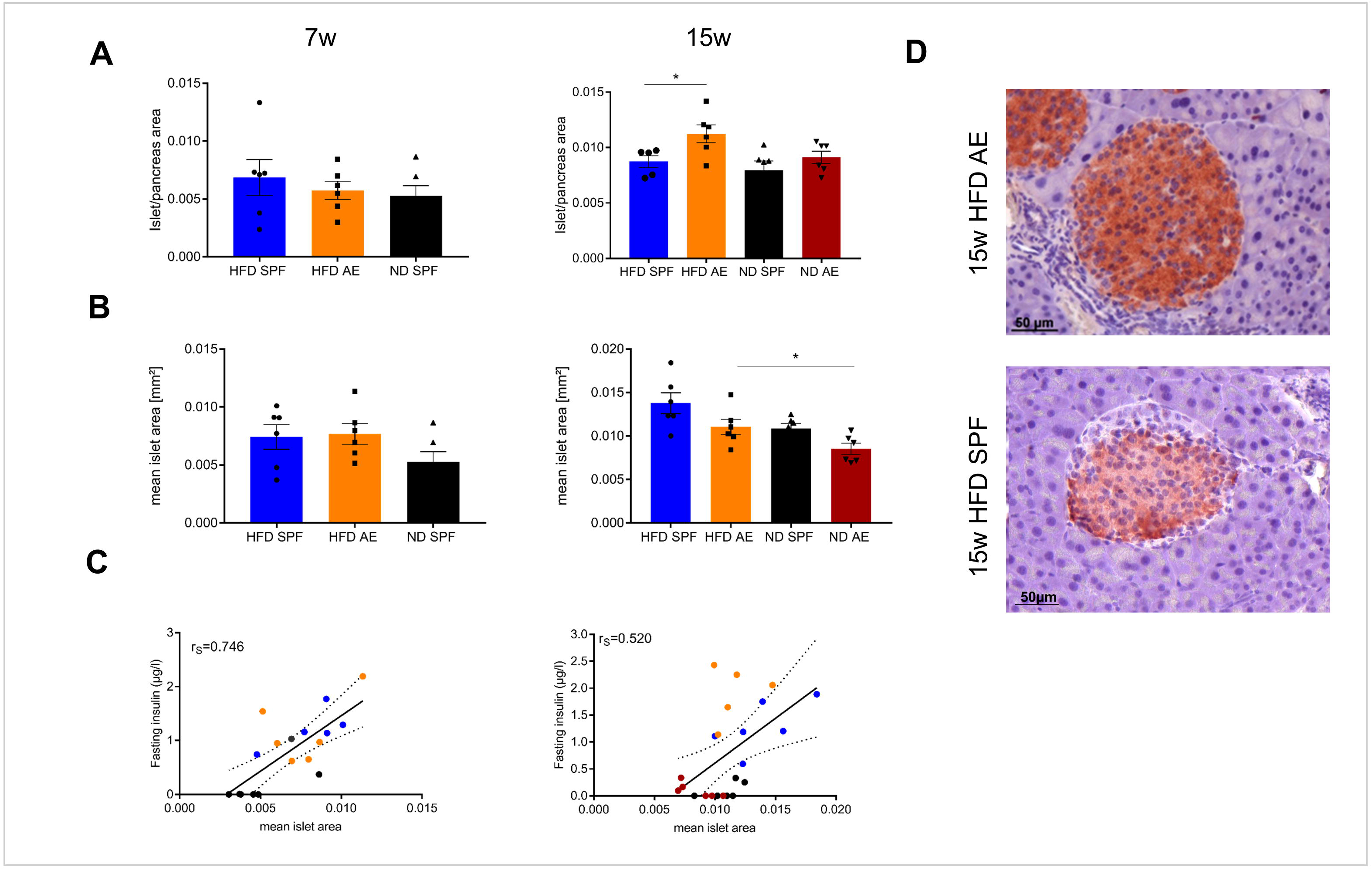
Antigen exposure protects pancreas morphology and function allowing lasting compensation of insulin resistance. Haematoxylin/eosin and immunohistochemical (IHC) insulin stainings were performed in 7 weeks and 15 weeks fed mice. **(A)** Mean islet cross-sectional area expressed as the percentage of the total pancreas area in 7weeks and 15weeks fed SPF and AE mice. Grubbs’analysis identified one outlier in 15weeks HFD fed SPF mice. **(B)** Mean islet area in mm^2^ for 7weeks and 15weeks fed SPF and AE mice. n=5-6 mice per group. **(C)** Mean islet area initially correlates significantly with fasting insulin in all groups (*P*=0.0004 (left), but correlation decreases over time because fasting insulin levels are kept high in AE mice only, *P*=0.011 (right). **(D)** Representative serial sections from mouse pancreas immunostained for insulin. Results are representative of 5-6 mice. Significance was determined using unpaired 2-sided Mann-Whitney U-Test or 1-way ANOVA. Scale bar, 50µm.

To sum up, histological analyses of pancreatic islets confirmed increased ß-cell compensation in AE mice that persisted over the observation time of 15 weeks HFD feeding.

### HFD in immune aged mice is a high risk combination for the development of NASH

Obesity is frequently associated with liver steatosis that can progress to NASH including a negative predictive value for the course of T2D and metabolic syndrome. Therefore, we next asked, whether housing conditions would have an impact on liver pathology. Intrahepatic naïve CD4^+^ and CD8^+^ T cells were found to be considerably lower in AE compared to SPF mice at week 7 (Fig. 7A, B). Conversely, for CD8^+^ T cells the fraction of effector memory cells of leucocytes was significantly higher (Fig. 7B), which was most pronounced in AE mice on HFD. Notably, more than 95% and 85% of CD4^+^ and CD8^+^ T cells, respectively, expressed the effector memory T cell phenotype in AE mice on HFD, a delta of > 25% compared to SPF mice on HFD (Fig. 7A-C). The innate cell subsets were less dramatically changed with trends for partial replacement of macrophages by infiltrating monocytes in AE mice and a slightly enhanced frequency of NK cells in both HFD groups (Fig. 7D, E). In addition, HFD mice displayed higher MHCII antigen expression levels under SPF conditions (Fig. 7F).

**Figure 7.**
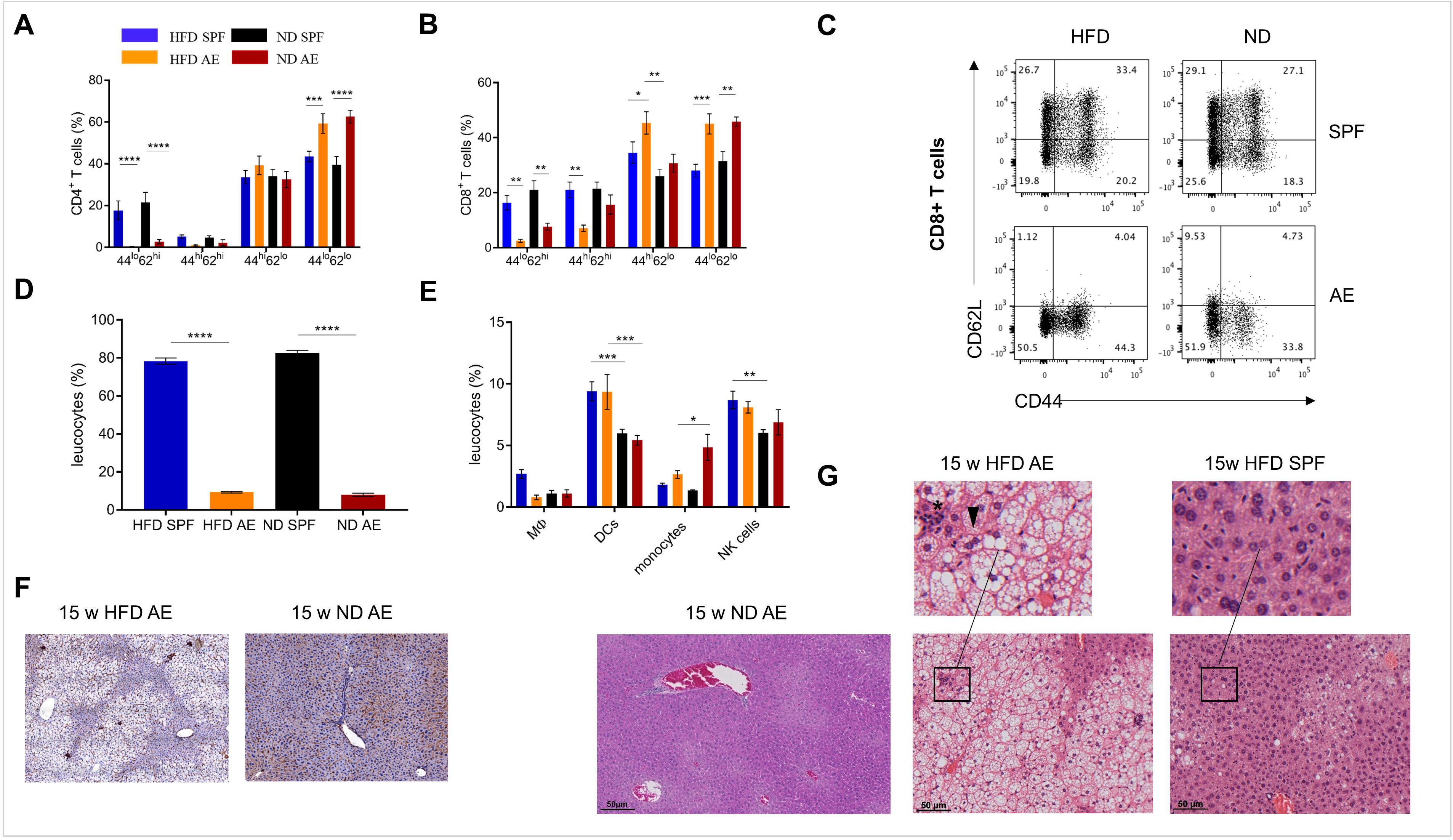
HFD in immune aged mice is a high risk combination for the development of NASH. Immune cell composition in livers of 7 weeks fed SPF and AE mice was analyzed via flow cytometry. **(A), (B), (C)** CD4^+^ and CD8^+^ T-cell liver phenotypes were compared among 7 week HFD and ND mice maintained under SPF and AE conditions. MHCII staining was performed in 15 weeks fed AE mice. **(D)**,**(F)** Percentages of mature (MHCII^high^) and immature (MHC^low^) dendritic cells (DCs) in liver tissue and representative IHC staining with MHCII in 15 week HFD and ND AE mice. **(E)** Percentages of innate immune cells in liver tissue. Haematoxylin/Eosin stainings were performed in 15 weeks fed SPF and AE mice. **(G)** Representative staining of ND AE and HFD SPF and AE mice. Infiltration of immune cells (asterix) and ballooned hepatocytes (arrowhead) illustrate NASH. n=5-10 mice per group. Significance was determined using 2-way ANOVA multiple measurement. *P<0.05, ** P<0.01, *** P<0.001, **** P<0.0001.

The liver weight did not differ between AE/SPF and HFD/ND mice (Supplementary Fig. 3B). However, HE staining of liver sections of mice on HFD for 15 weeks revealed severe steatosis in AE mice while only some SPF mice displayed a mild fat accumulation in the liver. In addition to a strong increase of large lipid droplets resulting in macrovesicular steatosis, the livers of AE mice on HFD showed lobular inflammation, hepatocyte injury in the form of hepatocellular ballooning and destroyed lobule structure. However, livers of ND AE mice did not display any signs of NASH (Fig. 7G). Sirius Red and Masson’s Trichrome staining revealed mild pericellular fibrosis in those mice (Supplementary Fig. 3C). Blinded NASH scoring of liver histology confirmed that 8/8 AE mice on HFD for 15 weeks displayed a strong NASH phenotype (score >3-5) whereas NASH was not observed in the majority of SPF mice (5/8, score <1) and the remaining liver sections were categorized as displaying only a mild form of NASH (3/8, score =3) (Supplementary Fig. 3A).

## Discussion

The data of this study confirm our hypothesis that “aging” of the adaptive immune system has a major impact on the course of obesity-related insulin resistance. Whereas obesity-induced insulin resistance is rather comparable in both groups fed a HFD, AE mice showed improved glucose tolerance by preserved compensatory ß-cell function (protective effect). In contrast, immunoaging was associated with rapid development of NASH (detrimental effect). The data underline the need for novel preclinical models that are closer to the human situation.

Worldwide, prevalence of obesity T2D is increasing (*23*). It is widely accepted that adipose tissue inflammation contributes to the development of obesity-related insulin resistance (*24, 25*). Various components of both the innate and the adaptive immune systems were identified as major players in regulating inflammatory processes in the development of insulin resistance (*26-28*), vasculitis and remodeling of parenchymal organs (*29-31*). However, knowledge about the exact pathomechanisms driving the process across the checkpoints is limited. Recently, awareness arose about the limitations of commonly used animal models regarding the shortcomings of clinical challenges (*32, 33*). With increasing knowledge about the role of the adaptive immune system, in particular of tissue-resident and tissue-infiltrating T cells, in controlling challenged tissue homeostasis it is more evident than ever that our widely used SPF mouse models do not reflect the “aging” immune system as seen in adults (*15, 16*).

Abnormally hygienic housing conditions of laboratory mice may account for limited translational potential to humans. To tackle this issue, we applied our recently developed model of AE housing and compared the development of insulin resistance, glucose tolerance, ß-cell function, and liver histology in two HFD fed groups: one kept SPF and the other outside the SPF barrier in AE conditions as described above.

Consistently, AE mice expressed a higher proportion of antigen experienced CD4^+^ and CD8^+^ T cells. In particular, memory effector T cell subsets were increased. Notably, AE mice did not acquire infectious diseases, as monitored following FELASA guidelines on a regular basis.

Our results demonstrate that changes in the immune cell composition of HFD fed C57Bl/6J mice affect the time course of glucose intolerance and insulin secretion. Interestingly, in contrast to our hypothesis, development of obesity-induced insulin resistance was comparable between the two HFD groups. This is in line with comparable immune cell subsets in visceral adipose tissue. Even in SPF mice, we observed a dominance of effector/memory T cells in visceral adipose tissue. Remarkably, HFD in both cohorts (SPF and AE) enhanced the effector/regulatory T cell ratio resulting in an immune disbalance in adipose tissue.

Intriguingly, AE mice fed a HFD revealed higher ß-cell responsiveness that was observed as an excess in insulin levels, which compensates for glucose intolerance. Furthermore, hyperinsulinemia in AE mice was in accordance with elevated body weight in comparison with SPF mice. As expected, pancreas histology revealed enlarged islet areas that correlated initially with fasting insulin levels in HFD mice. In contrast, ß-cells in SPF mice lost their functionality over time, confirmed by decreased insulin secretion and the strong correlation between islet mass and fasting insulin levels. Remarkably, ß-cells from AE mice were protected allowing continuous hyperinsulinemia. So far, the mechanisms behind the protective effects are unclear. The number of pancreas infiltrating T cells was roughly equal in both groups. Most T cells stuck to the perivascular region and did not enter the islets. As the pancreas expresses many IL-22 receptors (*34*), it could be speculated that Th17/22 cells might enter to secrete the pro-regenerative, islet-protective cytokine IL-22.

An alternative explanation might be that a vagus and/or sympathetic activation caused by systemic and local inflammation triggers the so-called “anti-inflammatory reflex” in pancreatic islets in AE mice, as described recently in critically ill patients suffering from major surgery or stroke-induced immunodeficiency (*35-37*).

Linked to the increasing prevalence of obesity worldwide, nonalcoholic steatohepatitis (NASH), the more inflammatory and progressive form of nonalcoholic fatty liver disease, emerges as a major health burden in developed countries (*38*). NASH is histologically characterized by ballooned hepatocytes, lipid accumulation, fibrosis and pericellular inflammation and may progress to cirrhosis, end stage liver disease or hepatocellular carcinoma (*39*). Several diets are known to induce NASH-like liver pathology in C57Bl/6J mice (*40*), but most of these relatively artificial approaches do not recapitulate human conditions of NASH and its metabolic consequences. Here, we show for the first time that mice maintained under non-SPF “antigen exposed” conditions develop NASH-like liver pathology as early as after 7 weeks of HFD feeding. They developed macrovesicular steatosis, hepatic infiltration and altered intrahepatic immune cell composition rarely seen in SPF mice even after 15 weeks on HFD. Recently, it has been described (*41*) that metabolic activation of intrahepatic immune cells causes NASH in C57Bl/6J mice in concordance with our results. Thus, our study offers a simple mouse model of short-term HFD for investigating the development of NASH and the underlying mechanisms that also exhibits fidelity to the human condition.

Based on our results, a number of questions arise. First, future studies are required to investigate the exact mechanisms of T cell activation in the liver and to tackle the question whether a systemic immune activation, induced by diet and housing conditions, or the dissemination of T cells activated in the liver accounts for NASH-like pathology and liver function. In contrast to these findings, severe liver inflammation was not accompanied by pancreatic tissue inflammation and did not correspond to ß-cell compensation for insulin resistance. Therefore, further studies have to be performed to validate the mechanisms of ß-protection, which are currently unknown. Similarly, in patients, severe steatohepatitis is often not attended by T2D, even though it is assumed that NASH is associated with insulin resistance (*42*). Second, the composition, diversity and function of the gut microbiome was described to be affected by HFDs (*43-45*), revealing the phenotypes seen in AE mice in a different light. Recently, it has been published that gut barrier function, e.g. intestinal permeability and alterations in intestinal levels of secondary bile acids, depend on housing conditions (*46*). The question whether gut colonization plays a causal role in adipose tissue and liver inflammation will be addressed in a forthcoming publication.

In conclusion, our study identifies an important new immunological link between environmental conditions and the progress of obesity, insulin resistance and its comorbidities. We provide a mouse model valuable for revealing new biomarkers of metabolic disease progression as well as testing novel therapeutic approaches.

## Acknowledgments

We thank Anke Jurisch, Diana Woellner, Kathrin Witte, Cornelia Heckmann, Christiane Gras and Francesca Liersch for assistance with experimental procedures and Benjamin Tiburzy from Biolegend for helpful comments on the gating strategy. Furthermore, we thank Mario Thiele for assistance with pancreas sections.

## Author Contributions Statement

J.S.-K. planned the study, wrote the manuscript, performed experiments and analyzed data, J.K. did immunostaining of liver and pancreas sections, S.B. helped with metabolic experiments, M.S. contributed to the selection of antibody panels and provided help with flow cytometry analysis, M.v.-H. contributed to conception and design of the study, A.K. performed CD3 staining of pancreas sections, K.S.-B. contributed to the invention of the AE mouse model, K.M. planned and carried out the clinical study “Effects of negative energy balance on muscle mass regulation” (registered at https://clinicaltrials.gov, NCT01105143), J.S. planned the concept of the study as endocrinological supervisor and reviewed the manuscript and H.-D.V. planned the concept of the study as immunological supervisor and wrote parts of the manuscript. J.S.-K., J.S. and H.-D.V. are the guarantors of this work and, as such, had full access to all the data in the study and take responsibility for the integrity of the data and the accuracy of the data analysis.

