## Supplementary Materials for "Distinct housing conditions reveal a major impact of adaptive immunity on the course of obesity-induced type 2 diabetes"

**Supplementary Table 1.** Housing conditions in specific pathogen-free (SPF) and antigen-exposed (AE) facilities. Animals in both facilities were monitored following FELASA guidelines.

**Supplementary Figure 1.** Wildtype SPF mice fed a HFD for 7 or 15 weeks develop glucose intolerance and insulin resistance. (A) Weight development in male 5 week-old C57BL/6 mice (n=10 per group) upon 7 or 15 weeks of HFD compared to mice fed ND. (B), (C) Quantitative analysis of the body composition by MRI and calculated weight of epididymal fat pads in g. (D) HOMA-index calculated as fasting plasma insulin (in milliunits per liter) × fasting plasma glucose (in mg per deciliter)/450. (E), (F) Blood glucose and insulin levels in an intraperitoneal glucose tolerance test (IPGTT) performed with 6 weeks and 12 weeks fed mice. (G) Respiratory Quotient calculated for 11 weeks fed mice during a 48h lasting observation period in metabolic cages. Significance was determined using 2-way ANOVA multiple measurement test. * P<0.05 (HFD 7week vs. ND 7week), + P<0.05 (HFD 15week vs. ND 15week), ** P<0.01 ((HFD 7week vs. ND 7week), ++P<0.01 (HFD 15week vs. ND 15week), *** P<0.001 (HFD 7week vs. ND 7week), +++ P<0.001 (HFD 15week vs. ND 15week), **** P<0.0001 (HFD 7week vs. ND 7week), ++++ P<0.0001 (HFD 15week vs. ND 15week).

**Supplementary Figure 2.** Diet-induced obesity rather than housing conditions change the immune cell composition in visceral adipose tissue. Immune cell composition was determined in visceral adipose tissue via flow cytometry in 7 weeks fed mice. **(A)** Heatmap showing relative distribution of T cell subsets and cells of the innate immune system as percentages of leucocytes in 7 week groups. Red indicates higher proportion, blue indicates lower proportion.

**(B)** Distribution of innate immune cells as percentage of leucocytes in HFD and ND mice maintained under SPF or AE conditions. **(C), (D)** Proportion of regulatory T and B cells in visceral adipose tissue of SPF or AE mice. **(E)** Percentages of CD44^lo^CD62L^hi^ (naϊve), CD44^hi^CD62L^hi^ (central memory), CD44^hi^CD62L^hi^ (effector memory) and CD44^lo^CD62L^lo^ (probably effector) T cells. Significance was determined using 2-way ANOVA multiple measurement **(B, E)** or ANOVA followed by Tukey´s multiple comparisons test **(C, D)** n=10 mice per group. *P<0.05, ** P<0.01, *** P<0.001, **** P<0.0001.

**Supplementary Figure 3 NASH-like liver pathology can be induced by HFD in AE mice.**

**(A)** Table showing histological scoring system for NASH. **(B)** Liver weight of 15weeks fed mice, n=10. Significance was determined using 1-way ANOVA * P<0.05. **(C)** Immunohistochemical staining of CD3-expressing T cells (upper row) and Sirius Red staining (lower row) in paraffin-embedded liver sections from mice fed an HFD or ND for 15 weeks. Arrows indicate CD3 staining, n=6-10mice.

**Supplementary Table 2.** Metabolic measurements correlate with immune cell subtypes in visceral adipose tissue of 15 weeks fed HFD SPF (left column) and AE (right column) mice. Significance was determined applying Spearman Correlation with 95% confidence interval. *P<0.05, ** P<0.01.

**Supplementary Table 3.** Antibodies used in flow cytometry staining. NIR-conjugated Zombie was used as live/dead marker.
