## Supplementary figures and images for "Distinct housing conditions reveal a major impact of adaptive immunity on the course of obesity-induced type 2 diabetes"

### Supplementary Materials

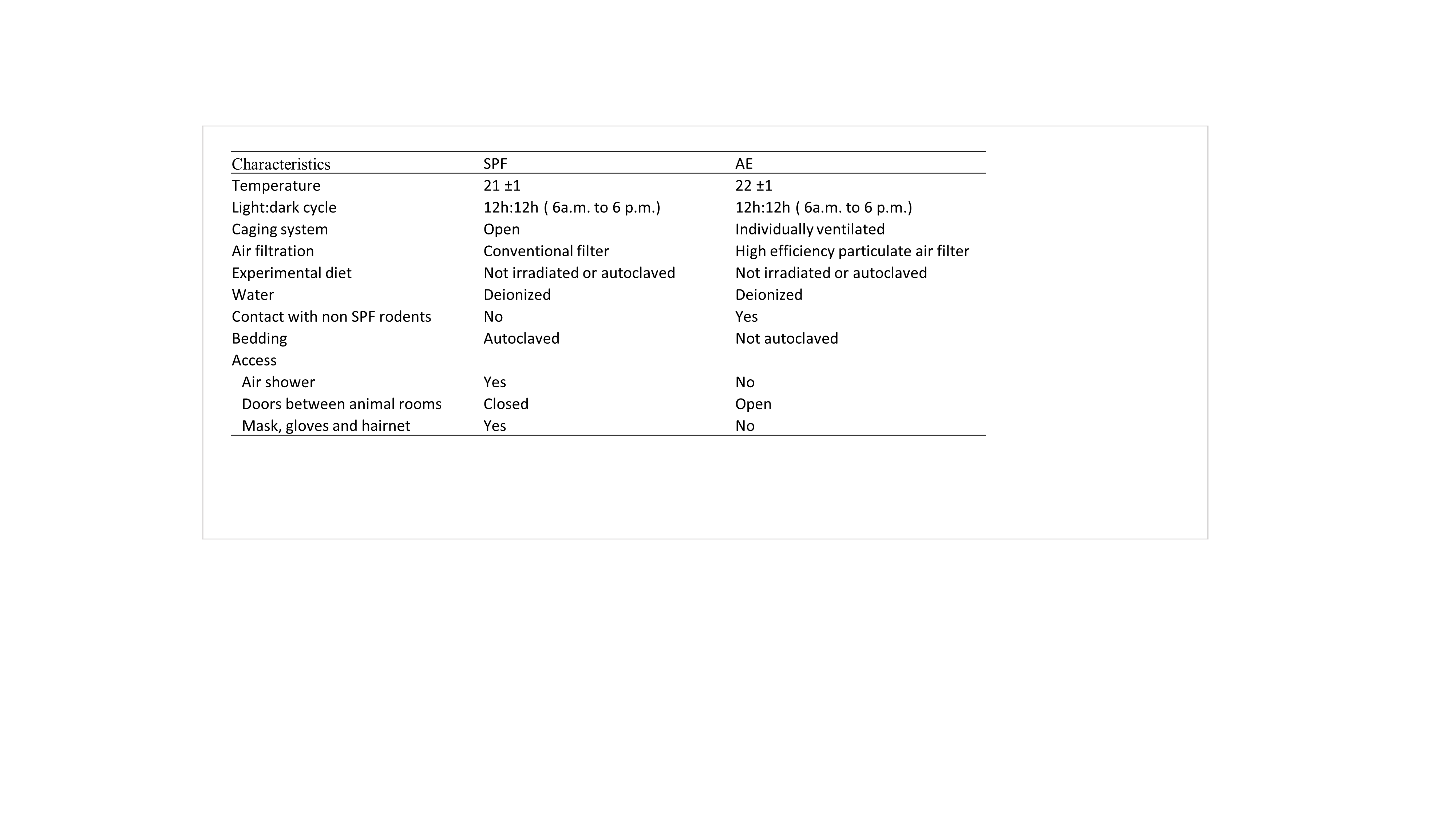

### Supplementary Materials

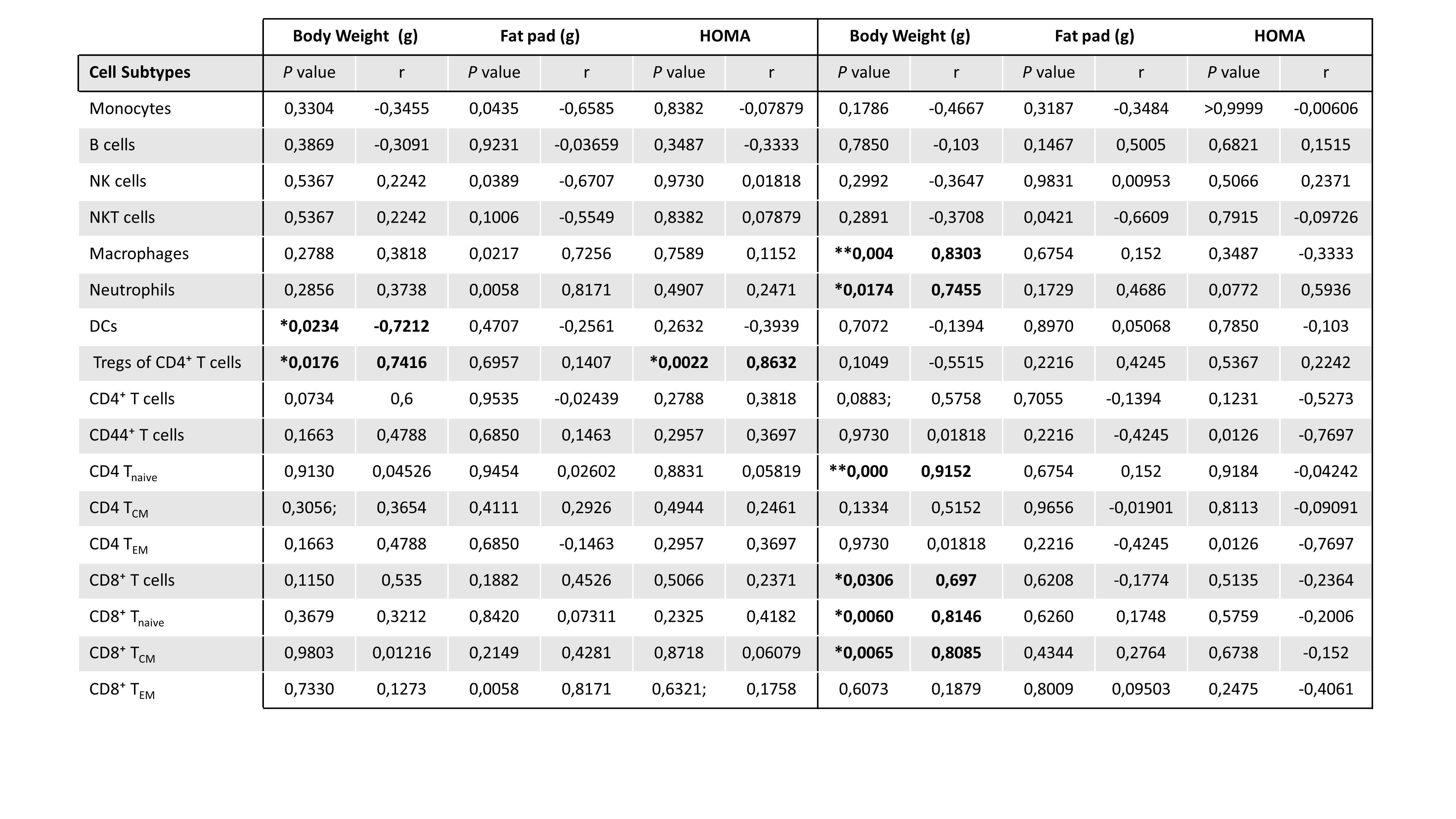

### Supplementary Materials

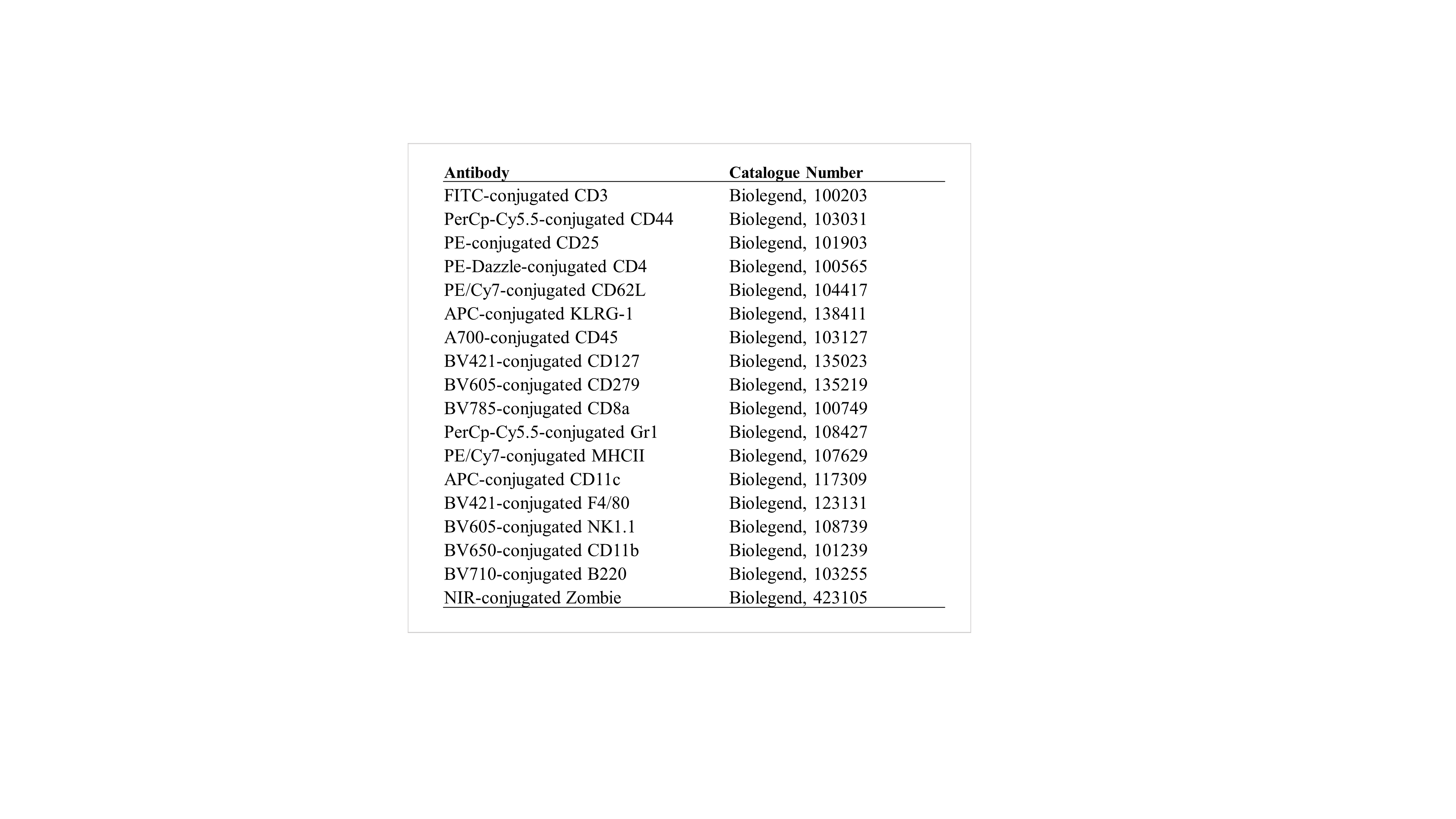

### Supplementary Materials

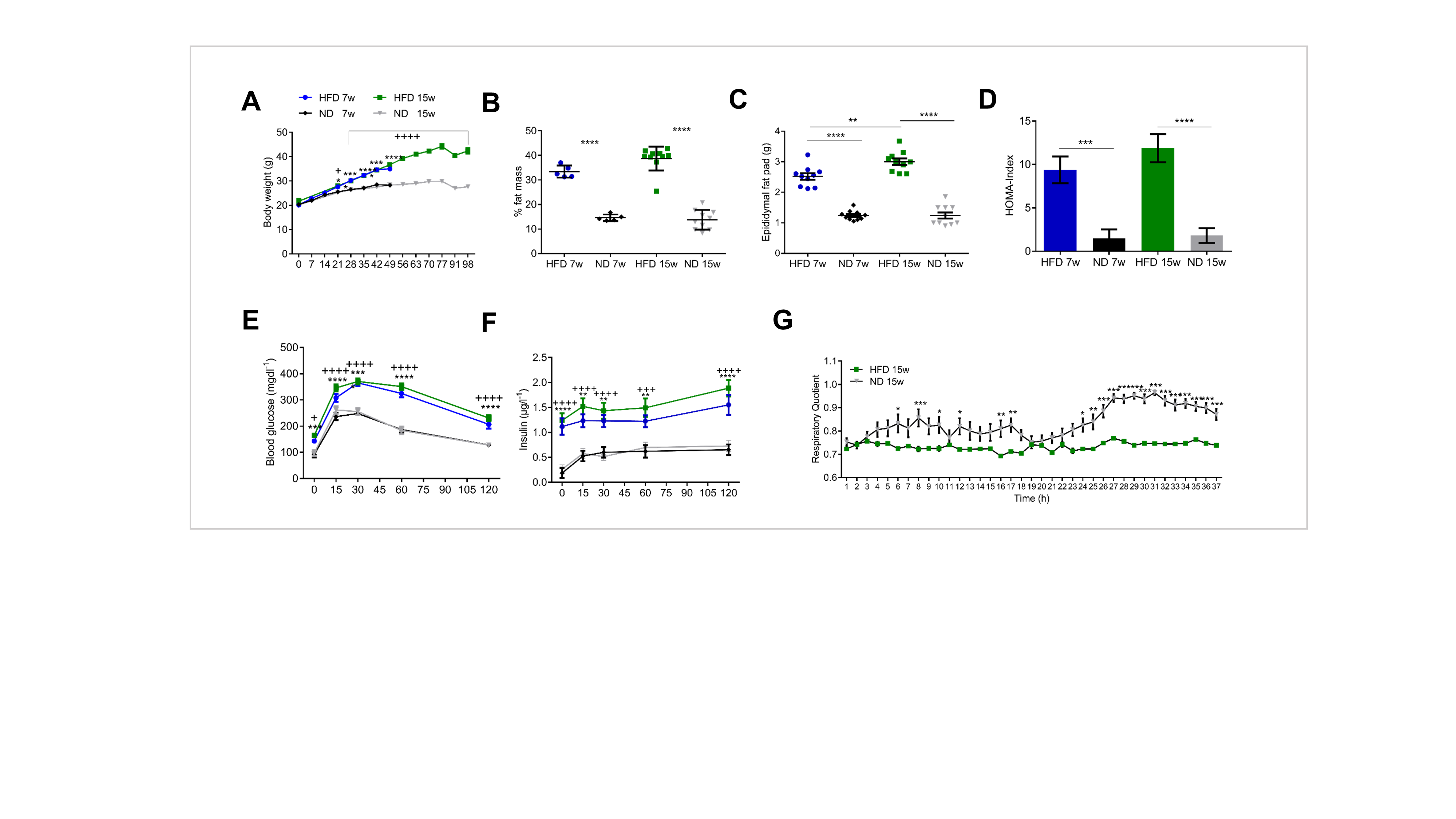

### Supplementary Materials

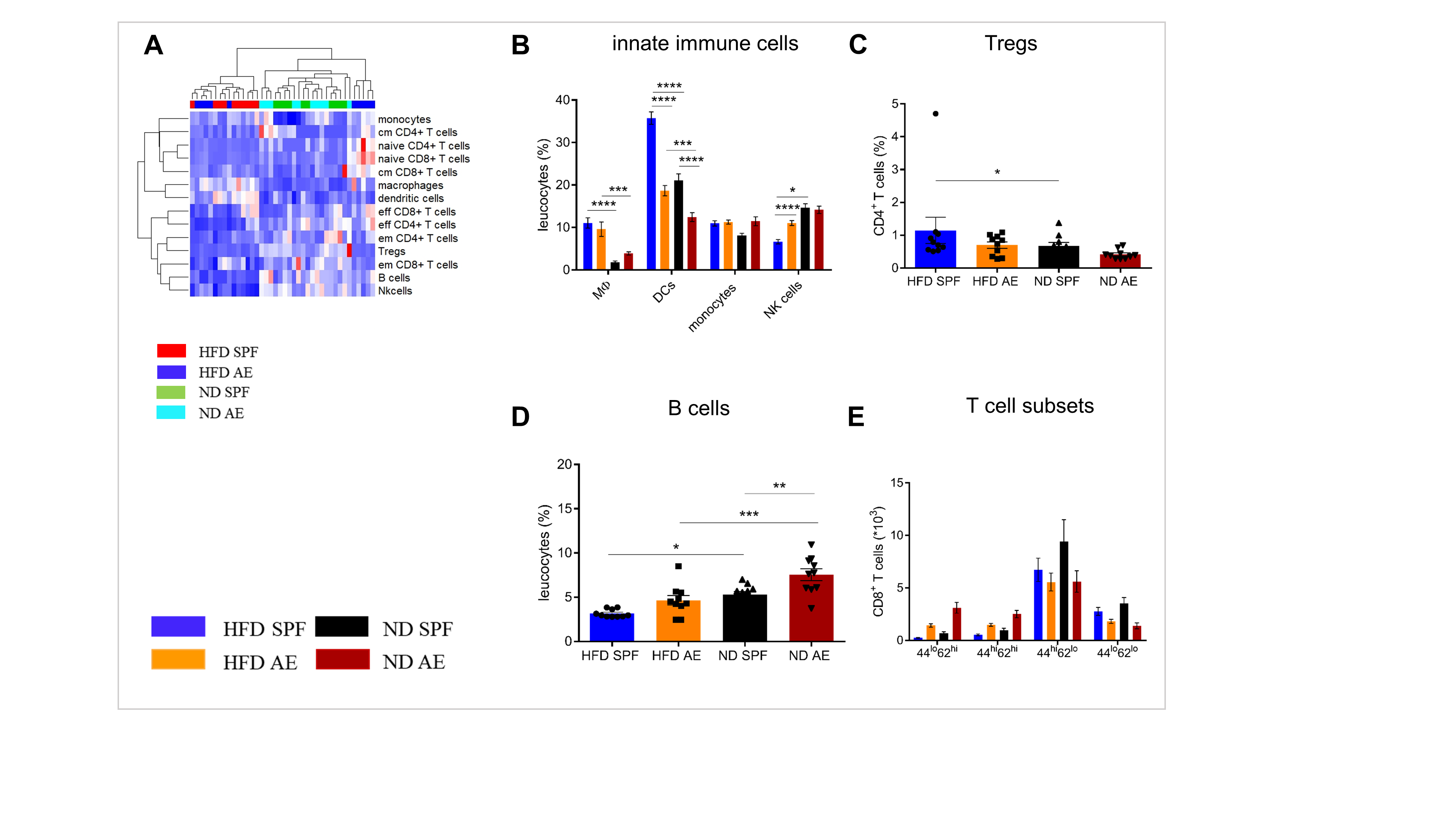

### Supplementary Materials

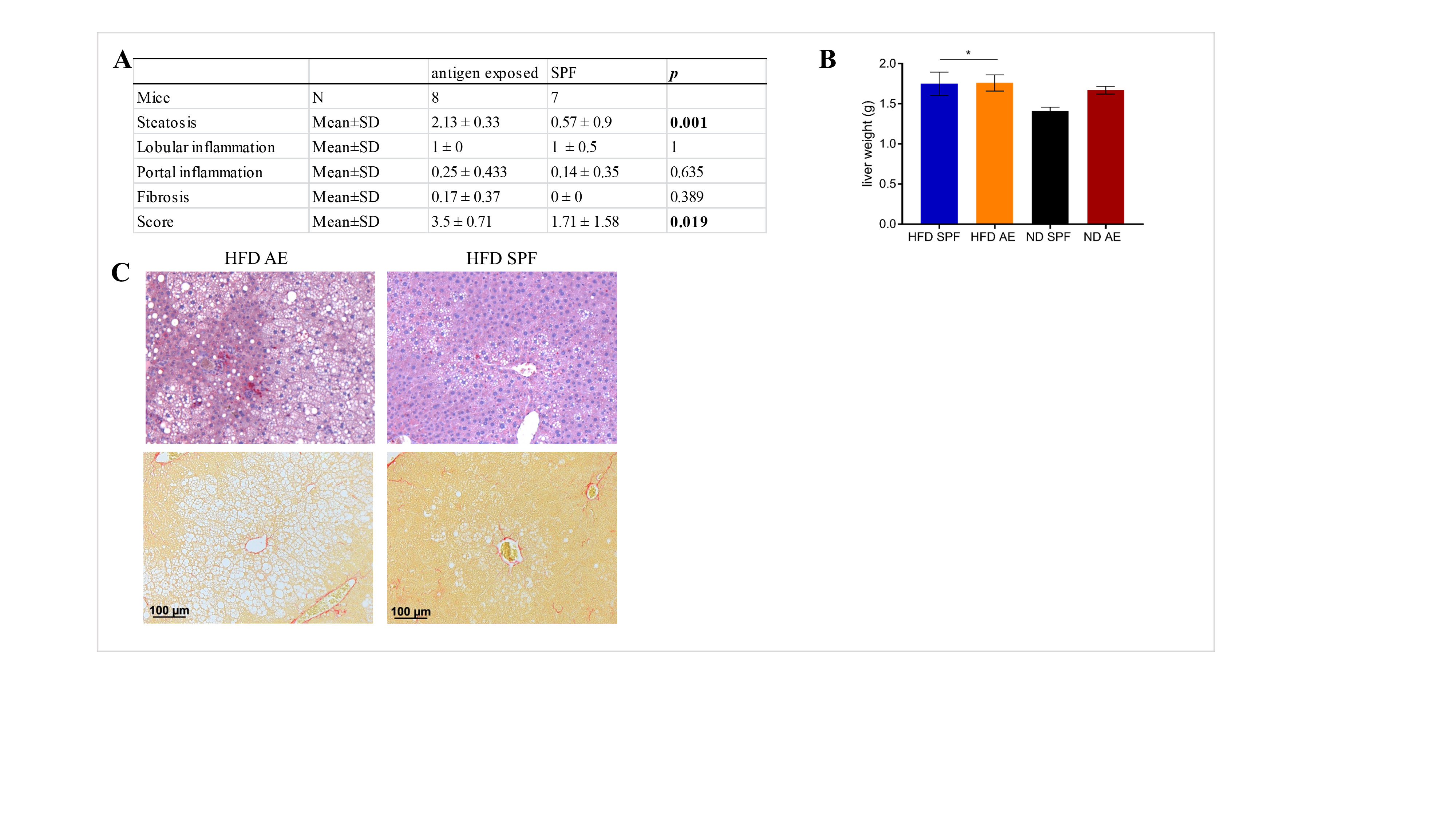
